# Does the territorial behaviour of the Amur tiger affect the accuracy of occupancy estimation?

**DOI:** 10.1101/2023.01.16.524338

**Authors:** Bing Xie, Yinan Gong, Yanwen Fu, Limin Feng, Haitao Yang

## Abstract

1. Accurate estimates of wildlife distributions and population persistence are essential for conservation programs. Occupancy models that account for detection probability have been used to characterize the occupancy and habitat selection of imperilled species. However, failure to distinguish between true-presence and pseudo-presence associated with territorial behaviour can result in the overestimation of the occupancy probability of target species in unsuitable habitats, and this can have major implications for the development of conservation strategies.

2. For highly territorial wildlife species requiring high-quality habitat for survival, occasional ‘Presence’ in unsuitable areas might be related to dispersal or migration, but this does not reflect actual occupancy and habitat use. ‘True-Presence’ and ‘Pseudo-Presence’ should be distinguished for target species to better reflect their actual occupancy and habitat use.

3. To investigate the effect of ‘True-Presence’ and ‘Pseudo-Presence’ on wildlife occupancy estimation, we developed a modified model (M_m_) that considers the territorial behaviour of the Amur tiger in northeast China, which distinguished between ‘True-Presence’ and ‘Pseudo-Presence’. We compared two models, M_m_ and M_0_ (conventional occupancy model), and assessed model performance using goodness-of-fit evaluation, detection and occupancy probability, and favourable variable selection.

4. We found that M_m_, which has fewer favourable variables, is more powerful than M_0_ for estimating detection and occupancy probability, as well as characterizing the effect of various factors on the habitat use of Amur tigers. Furthermore, M_m_ significantly reduced the overestimation of occupancy probability outside the home range compared with M_0_, and detection probability estimates did not significantly differ between M_0_ and M_m_. Finally, M_m_ provided more empirical habitat selection variables for the Amur tiger.

5. Our results revealed how ‘True-Presence’ and ‘Pseudo-Presence’ affect the occupancy probability and habitat selection of Amur tigers. Our method improves the accuracy of occupancy models; it can also be used to characterize the habitat selection and distribution of wildlife species and be applied to other territorial species.

## Introduction

Occupancy is defined as the probability that a randomly selected site or sampling unit in an area of interest is occupied by a species (MacKenzie *et al*. 2002; MacKenzie *et al*. 2017). Studies of the occupancy of wildlife typically evaluate the effects of various environmental factors on the habitat selection and distribution of wildlife species. Occupancy models that account for detection probability can provide more robust estimates of occupancy as well as the effects of various factors on occupancy (MacKenzie *et al*. 2002; MacKenzie *et al*. 2003; MacKenzie *et al*. 2005). Occupancy models have been used to model several ecological issues, including species distributions (Kéry, Guillera-Arroita & Lahoz-Monfort 2013), interactions between species and habitats (Karanth *et al*. 2011), dynamics of mate-population (Ferraz *et al*. 2007), and interspecific relationships (Miller *et al*. 2012) in multiple taxa, such as mammals (Yang *et al*. 2021), birds (Goodwin & Shriver 2011), amphibians (Tournier *et al*. 2017), plants (Chen *et al*. 2013) and fish (Falke *et al*. 2012).

The occupancy model is implemented by carrying out repeated surveys to obtain the ‘Presence-Absence’ history of the target species to minimize the probability of ‘false absences’ from a sample unit (MacKenzie *et al*. 2017). Based on MacKenzie *et al*. (2002), ‘Presence’ of target species in a sample unit indicates that a certain unit is by default occupied; however, the ‘Absence’ of target species in a sample unit indicates that the unit is uncertain and requires estimation from the model. Can the occupancy status or habitat use of target species be confirmed by the ‘Presence’? Does the occasional ‘Presence’ in a highly unsuitable area reflect the actual occupancy status or habitat use of the target species? For highly territorial wildlife species requiring high-quality habitats, the occasional ‘Presence’ in an unsuitable area might reflect behaviours such as dispersal or migration (Berigan *et al*. 2019), which do not reflect actual occupancy or habitat use. To better characterize the habitat use and distribution of wildlife species, ‘True-Presences’ (i.e., detections of the target species in an area that reflect actual occupancy or habitat use) and ‘Pseudo-Presences’ (i.e., detections of the target species in an area that do not reflect actual occupancy or habitat use) should be distinguished. Failure to distinguish between ‘True-Presence’ and ‘Pseudo-Presence’ can result in the overestimation of the occupancy probability of target species in unsuitable habitats, and this can have major implications for the development of conservation strategies.

Felidae, which are mostly solitary and require large tracts of high-quality habitat for survival, maintain home ranges that are not only necessary for meeting energetic demands but are also important for enhancing fitness and reproductive success (Sunquist & Sunquist 2017; Sandell 2019). The area of felids’ home range only changes dramatically when female recruits need area to reside or male recruits replace the local male residents (Smith 1984). The tracks of felids are seldom detected outside of their home range, given that detections outside of home ranges generally reflect dispersal or eviction. Thus, ‘True-Presence’ (i.e., detections inside home ranges) and ‘Pseudo-Presence’ (i.e., detections outside of home ranges) should be accounted for in occupancy models to ensure that accurate information regarding habitat use and population status is obtained.

A literature search on Google Scholar from 2002 to December 31, 2020, with the keywords ‘occupancy model’, ‘Felidae’, ‘Panthera’, and ‘habitat’ revealed 728 papers that have applied occupancy models to Felidae species. Ninety-seven papers on 29 felid species cited MacKenzie *et al*. (2002), which included 63 scientific papers, 32 theses, and 2 books (Fig. 1) (Appendix 1). The highest numbers of published papers were observed in 2019 and 2020 (both 16), which indicates that occupancy models have become an effective method for studying Felidae species. All these papers explored the problem of imperfect detection (detection probabilities < 1), but none of them have distinguished between ‘True-Presence’ and ‘Pseudo-Presence’ associated with the effect of felid behaviour on space use or habitat selection. Consequently, the area of the distribution estimated for several felid species in these previous studies might be overestimated.

**Figure 1.**
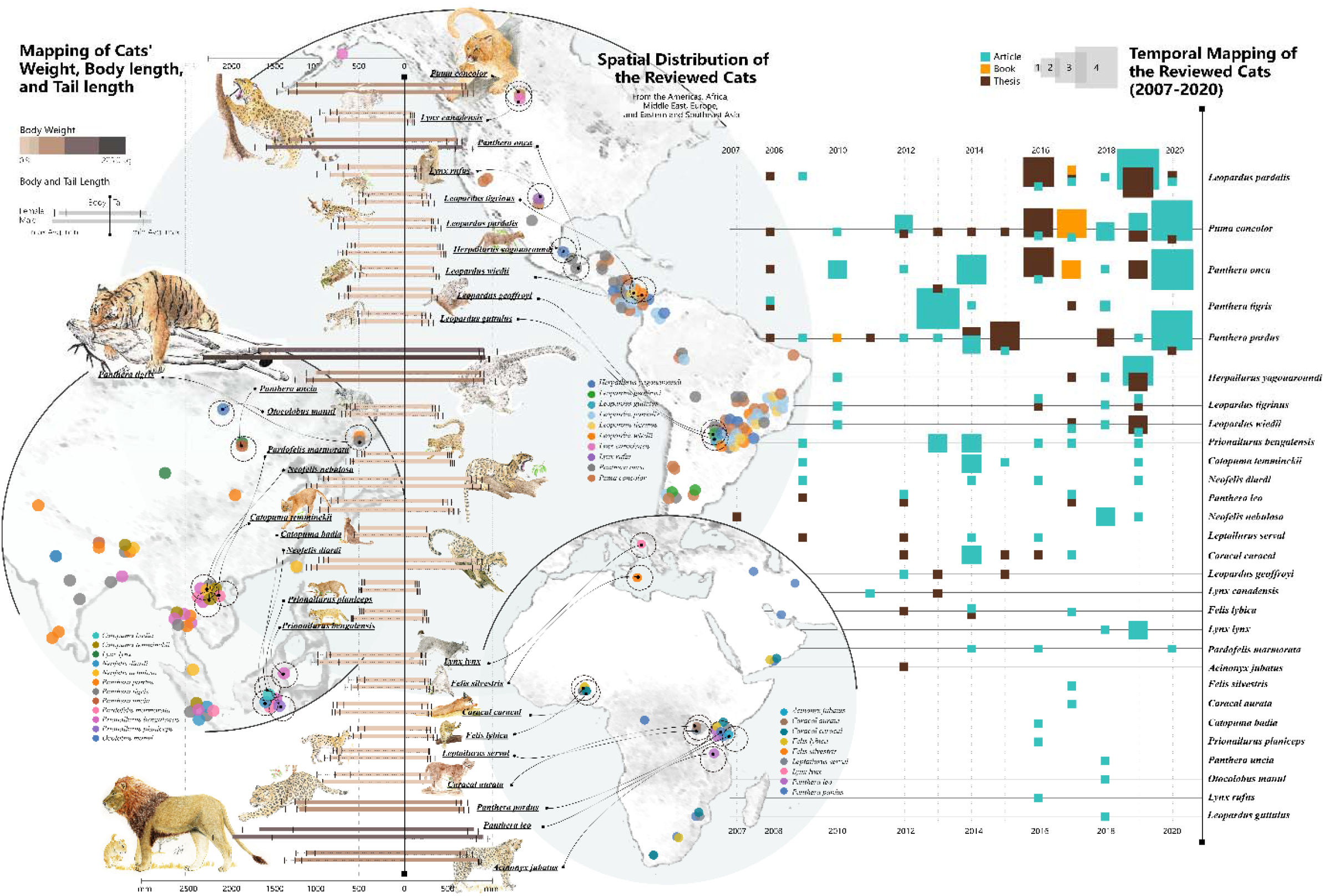
Published MacKenzie et al. (2002) occupancy method-based occupancy estimation in relation to Felidae species. The column “Mapping of Cats’ Weight, Body length, and Tail length” showed all 29 felids which cited MacKenzie et al. (2002); the column “Spatial Distribution of the Reviewed Cats” showed the geographic region of published studies; the column “Temporal Mapping of the Reviewed Cats” showed the published studies’ number of per cat species since 2002

In this study, we used occupancy models based on camera trapping data to study the distribution of the Amur tiger (*Panthera tigris altaica*) in Northeast China. We evaluated the performance of the conventional occupancy model (M_0_, which does not distinguish between ‘True-Presence’ and ‘Pseudo-Presence’) and the modified occupancy model (M_m_, which distinguishes between ‘True-Presence’ and ‘Pseudo-Presence’). Our objectives were twofold. First, we designed a practical M_m_ that considers the territorial behaviour of Amur tigers to generate reliable estimates of their distribution. Second, we evaluated the effect of environmental factors on the habitat of the Amur tiger to verify the accuracy of M_m_. We hypothesized that 1) the detection probabilities of the Amur tiger would not significantly differ between M_0_ and M_m_, and 2) the occupancy probabilities of the Amur tiger would significantly differ between M_0_ and M_m_. We used the occupancy models of the Amur tiger in this study as a case to explore the effect of the ‘True-Presence’ and ‘Pseudo-Presence’ on the occupancy models, especially occupancy models, which were applied to estimate habitat use of Felidae at a fine scale.

## Materials and methods

### Study area

We conducted camera trapping surveys in Northeast Tiger and Leopard National Park (NTLNP) in northeastern China and the Land of the Leopard National Park (LLNP) in Russia’s southwest Primorye Province (Fig. 2). Transboundary collaboration between China and Russia on the Amur tiger and Amur leopard has greatly enhanced the monitoring, research, and conservation of these endangered felid species (Vitkalova *et al*. 2018). Most tigers and leopards occur near the border (Wang *et al*. 2016; Wang *et al*. 2017; Vitkalova *et al*. 2018). The Amur tigers and leopards can move freely across the border and establish home ranges that span both countries.

**Figure 2.**
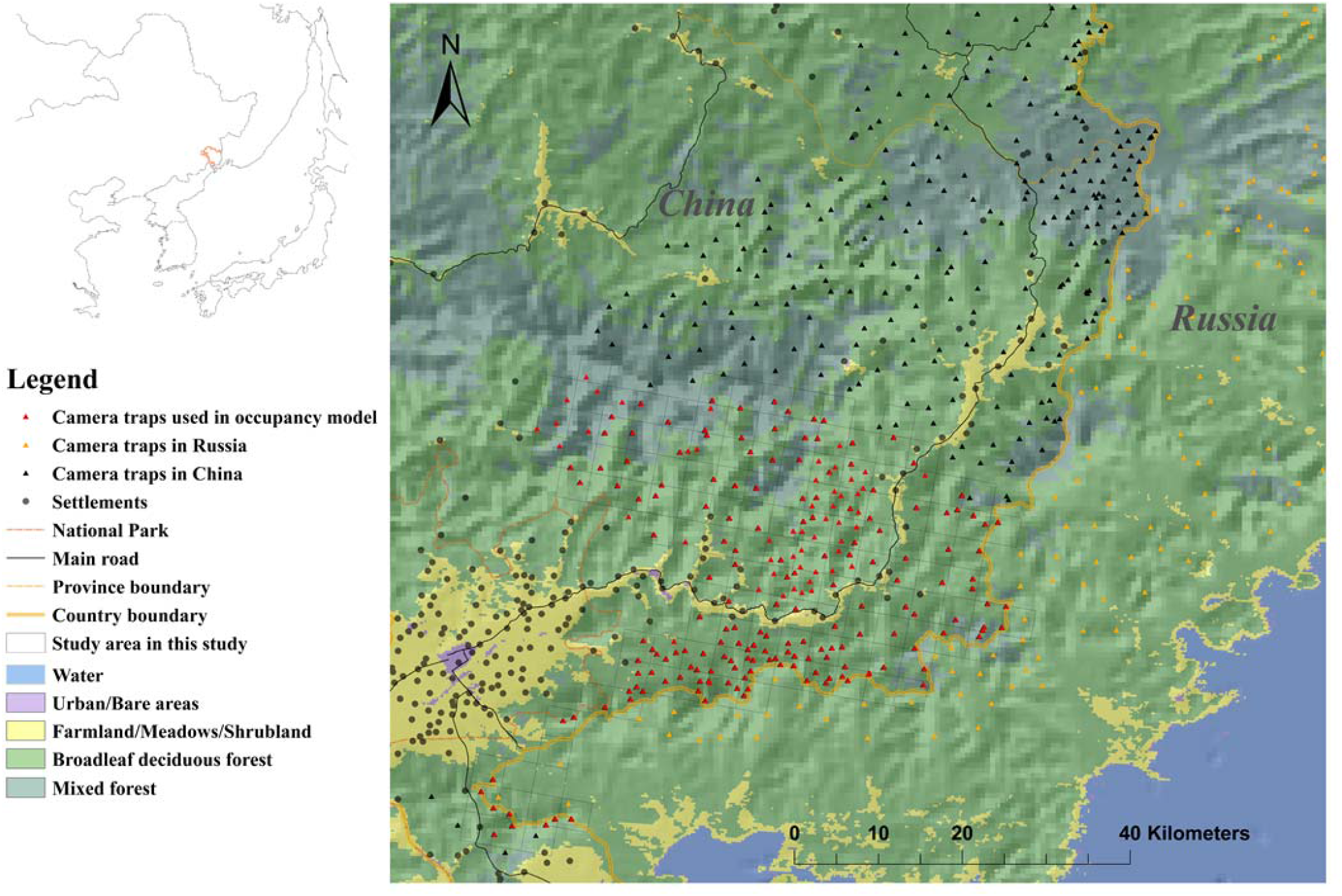
Map of the area sampled by camera traps in China and Russia, showing the camera sites used in this study and relative to settlements, roads and different land cover. The inset shows the location of the study area within China.

We can precisely estimate the home ranges of individual tigers based on camera trapping images, including the home ranges of individuals that have been sighted in both China and Russia. Our analyses were based on reliable estimates of the home ranges of Amur tigers. The data used in the following occupancy model in this study were for individuals with home ranges that were exclusively in China. For detailed climatic and geographic information, see Appendix 2 *Study area*.

### Data collection

In China, the study area was divided into a 3.6 × 3.6-km grid, and 1–4 camera trapping sites (mostly double-sided) were established in each grid cell, with the exception of those without forests (farmland and villages). In Russia, the study area was divided into a 3 × 3-km grid, and 1–2 camera trapping sites were established in each cell (Fig. 2) (Vitkalova & Shevtsova 2016). Cameras were placed on trees at a height of 0.4 to 0.8 m above the ground, along ridges, forest roads, and paths frequented by tigers and prey species (Wild boar *Sus scrofa*, Siberian roe deer *Capreolus pygargus*, and Sika deer *Cervus nippon*) as well as humans. There were a total of 218 camera trapping sites in China and 143 camera trapping sites in Russia (Fig. 2) (Vitkalova *et al*. 2018). We identified individual tigers based on their distinct skin patterns (Karanth & Nichols 1998), and sex was determined by the presence or absence of testicles. To avoid inflated counts caused by repeated detections of the same event, we excluded consecutive recordings of the same species within 0.5 h (O’Brien, Kinnaird & Wibisono 2003). The relative abundance index (RAI: the number of detections/100 trap days) was calculated for the prey species and human activity.

### Modified occupancy model

Female tigers only breed when they occupy a home range, and the number of adult females determines the recruitment potential of the tiger population (Smith 1987). The home ranges of male tigers can change rapidly, and the stable home ranges of female tigers play a key role in ensuring the viability and resilience of populations, as well as determining the distribution of adult males (Smith 1984; Smith 1987; Goodrich *et al*. 2010; Simcharoen *et al*. 2014). Study of resident female tigers would provide a more accurate characterization of habitat selection. Thus, we focused exclusively on the home ranges of resident female tigers and treated adult males and non-resident tigers as an important variable (T00) in M_m_ (Table 1). We estimated the home ranges of female tigers using 100% minimum convex polygons (MCP) from June 2011 to April 2016 (Table S1) following the definition of resident tigers from Goodrich *et al*. (2010). Although outliers might affect the 100% MCP estimation and induce inflation (Harris *et al*. 1990; Simcharoen *et al*. 2014), we used MCP estimation because our sample sizes for determining the potential home range of each tiger were small (ranging from 18 to 28 camera trapping sites) as well as to facilitate comparison with other studies (Goodrich *et al*. 2010; Gil-Sánchez *et al*. 2011; Simcharoen *et al*. 2014).

**Table 1.** Variables considered for occupancy models to predict habitat use of the Amur tiger

| Variables | Description | Category | Source | Parameters |
| --- | --- | --- | --- | --- |
| Effort<sup>1, 2</sup> | Numeric, Number of days camera trap was active for each trap site | - | Camera trap | $P$ |
| Non-resided tigers and males<sup>2</sup> | Numeric, Relative Abundance Index (RAI)<sup>a</sup> of non-resided tigers and males (T00) | Habitat | Camera trap | $\psi$ |
| Home range<sup>2</sup> | Categorical, Located in home ranges of tiger individuals (1) or not (0) (HOME) | Habitat | Camera trap | $\psi$ |
| Forest type<sup>1,2</sup> | Categorical, Broad-leaved forest, Mixed forest, Coniferous and Oak forest<sup>b</sup> (FT) | Habitat | Field research | $P, \psi$ |
| Elevation<sup>1, 2</sup> | Numeric(m), Elevation of each camera-trap site (ELE) | Habitat | GPS record | $\psi$ |
| Sika deer<sup>1, 2</sup> | Numeric, Relative Abundance Index (RAI) of sika deer (S05) | Prey | Camera trap | $\psi$ |
| Roe deer<sup>1, 2</sup> | Numeric, Relative Abundance Index (RAI) of wild boar (S04) | Prey | Camera trap | $\psi$ |
| Wild boar<sup>1, 2</sup> | Numeric, Relative Abundance Index (RAI) of roe deer (S03) | Prey | Camera trap | $\psi$ |
| Human activity<sup>1, 2</sup> | Numeric, Relative Abundance Index (RAI) of human and vehicle (HUVe) | Disturbance | Camera trap | $\psi$ |
| Livestock<sup>1, 2</sup> | Numeric, Relative Abundance Index (RAI) of livestock (LIV) | Disturbance | Camera trap | $\psi$ |
| Road type <sup>1, 2</sup> | Categorical, Dirt road, Logging road, Ridge, Forest trail and Animal road (ROAD)<sup>c</sup> | Habitat | Field research | $P, \psi$ |
| Road distance <sup>1, 2</sup> | Numeric(m), Distance to the nearest main road from camera-trap site (RD) | Disturbance | Measured from google earth | $\psi$ |
| Settlement distance <sup>1, 2</sup> | Numeric(m), Distance to the nearest settlement from camera-trap site (SED) | Disturbance | Measured from google earth | $\psi$ |

NOTE: <sup>1</sup> Variables were used in $M_0$ ; <sup>2</sup> Variables were used in $M_m$ ; <sup>a</sup> Relative Abundance Index (RAI): independent detection frequency per 100 camera-trap days; <sup>b</sup> Broad-leaved forest: the dominant species of broad-leaved forest is not oak; Mixed forest: the dominant species is not only broad-leaved tree but also conifers; Coniferous: the dominant species is conifers; Oak forest: the dominant species is oak; <sup>c</sup> Dirt road: the main road in our study area used by vehicles; Logging road: the abandoned logging road for motorcycles or foot traffic; Ridge: trail on the ridge; Forest trail: the collection road for local people on foot and linked to the dirt road or logging road; Animal road: the road used by animals and local people and not linked to the dirt road or logging road. $P$ : probability of detection; $\psi$ : probability of occupancy.

We built and assessed two single-season occupancy models for M_0_ and M_m_ following the assumptions of MacKenzie *et al*. (2002). Camera trapping data were used in M_0_ and M_m_ from a 112-day window (July 1 to October 20, 2014), which was divided into eight occasions. The detection histories of tigers for each camera trapping site were developed based on records from the cameras, where “1” indicates that a tiger was detected at a specific camera trapping site on a specific occasion, and “0” indicates no detection. For M_0_, we counted all tiger occurrences as “1”, but only ‘True-Presence’ was considered “1” for M_m_ (Appendix 3).

We selected 11 variables for M_0_, including prey species, human activities, and environmental factors (Table 1) (see details in Yang *et al*. (2018) and Yang *et al*. (2019)). In addition, we created two unique variables for M_m_, non-resident tigers (T00) and home range (HOME). T00 is a numeric variable that indicates the relative abundance index (RAI) of non-resident tigers and male tigers at a given camera trapping site (Table 1). HOME is a categorical variable indicating whether the camera trapping site was located in the area of a female’s home range (100% MCP area) (“1”) or not (“0”) (Table 1). Before adding variables to the model, we checked for collinearity using the variance inflation factor (VIF) and correlation coefficient (r), excluding those with VIF > 3 or |r| > 0.7 (Yang *et al*. 2019).

Hierarchical occupancy models (*probit* mixture framework and reduced-dimensional spatial process) were used to improve algorithm convergence and eliminate spatial autocorrelation (Johnson *et al*. 2013), which ensured spatial independence (MacKenzie *et al*. 2002). We ran models in two phases (Yang *et al*. 2021): 1) models without spatial autocorrelation to select covariates using the *unmarked* package in R (Fiske & Chandler 2011); 2) Bayesian models incorporating the spatial random effect of the selected models using the *stocc* package in R (Johnson *et al*. 2013). For detailed information on the occupancy models, see Appendix 2 *Modified occupancy model*. Finally, we compared the occupancy and detection probabilities of M_0_ and M_m_ using Mann-Whitney *u* statistics.

## Results

### Home ranges of adult females

The home ranges of five resident female tigers were estimated, and the average home range size was 257.95 ± 74.84 km^2^ (mean ± SD) (Fig. 3, Table S1). We obtained the study area for the occupancy models, which included all home ranges of females and the area outside of these home ranges using MCP on the Chinese side (Fig. 2). The size of our study area was 1,750 km^2^, and there were 218 camera trapping sites (153 inside the home range area and 65 outside the home range area) in the study area (Figs. 2, 3).

**Figure 3.**
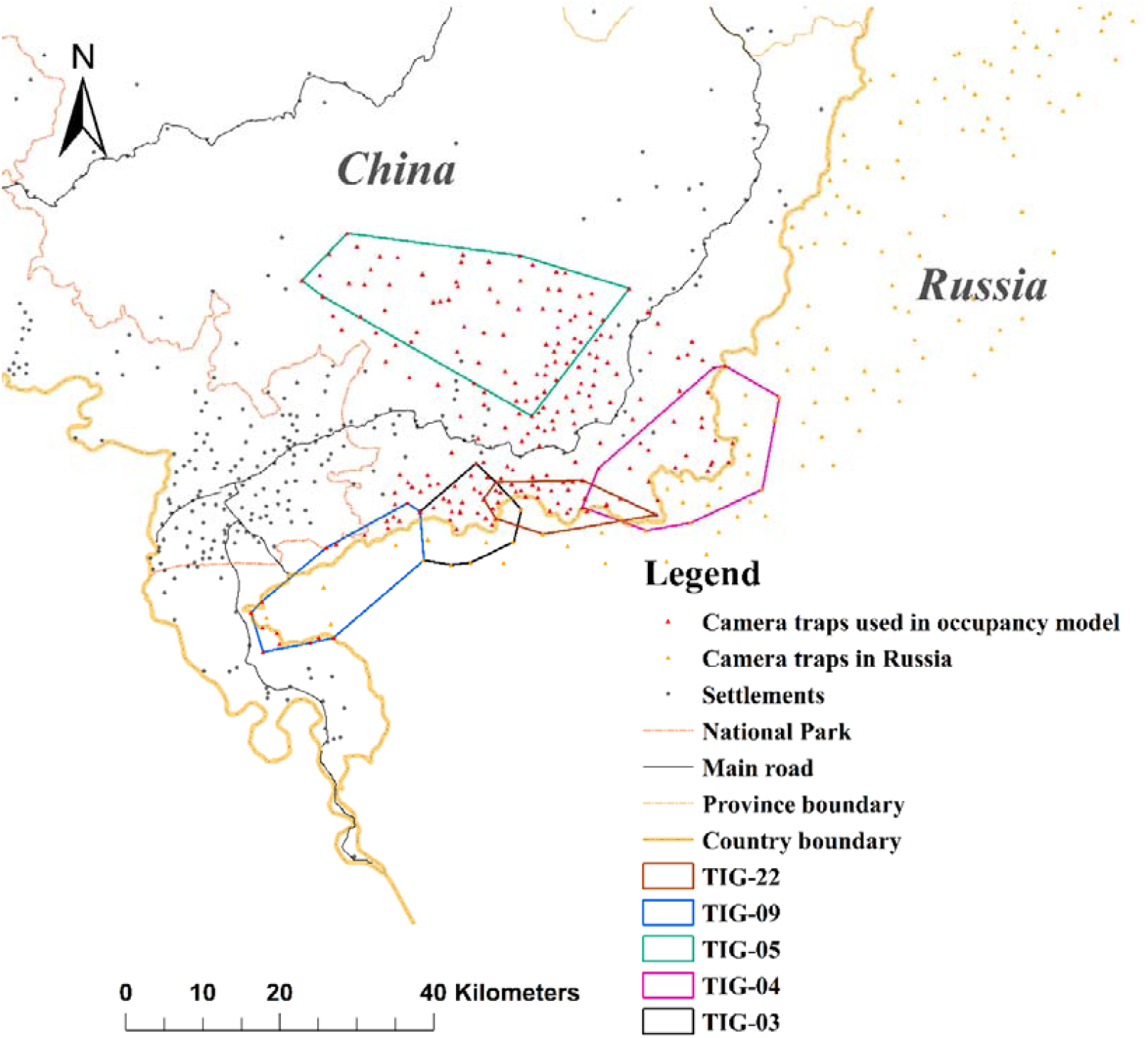
Map of the home ranges of 5 female tigers in the study period.

### Model performance

We recorded 425 detections of tigers at 218 camera trapping sites during the study period, which were active for 22,716 days (average effort was 104 ± 1.34 days (mean ± SE)). None of the variables were eliminated from any model (max VIF = 2.44 < 3, and |r| = 0.53 < 0.7) (Table S2).

Under the maximum likelihood framework, the top-ranked detection model (∆AIC < 2), *P*(ROAD)ψ(.), was used in M_0_ and M_m_ (Table 2, Table S3) for occupancy probability estimation. There were 11 and 2 top models for the occupancy estimations (∆AIC < 2) for M_0_ and M_m_, respectively (Table 2). Both M_0_ and M_m_ had high goodness-of-fit (M_0_: *P* = 0.79, c-hat = 0.89; M_m_: *P* = 0.57, c-hat = 0.96). Four and 6 variables for M_0_ and M_m_, respectively, with summed model weights > 0.5 were used to assess spatial autocorrelation (Table 2, Table S3). The results of the RSR models (model spatial autocorrelation with restricted spatial regression) indicated that the random spatial effects do not require consideration in M_0_; the goodness-of-fit values without spatial autocorrelation and the RSR models were almost equal in M_m_, which facilitates comparison with the modified occupancy models. We used the results of the non-RSR models for both M_0_ and M_m_ (Table S4).

**Table 2.** Top-ranked models (ΔAIC ≤ 2) for M_0_ and M_m_.

| Models | AIC | $\Delta AIC$ | AIC $W_t$ | Cum. $W_t$ |
| --- | --- | --- | --- | --- |
| $M_0$ | | | | |
| $P(ROAD)\psi(S05+HUVe+SED+S04)$ | 724.15 | 0.00 | 0.166 | 0.17 |
| $P(ROAD)\psi(S05+HUVe+SED)$ | 724.60 | 0.45 | 0.133 | 0.30 |
| $P(ROAD)\psi(S05+HUVe+SED+S03)$ | 725.39 | 1.24 | 0.089 | 0.39 |
| $P(ROAD)\psi(S05+HUVe+SED+ELE)$ | 725.42 | 1.26 | 0.088 | 0.48 |
| $P(ROAD)\psi(S05+HUVe+SED+S04+RD)$ | 725.56 | 1.40 | 0.082 | 0.56 |
| $P(ROAD)\psi(S05+HUVe)$ | 725.57 | 1.42 | 0.081 | 0.64 |
| $P(ROAD)\psi(S05+HUVe+SED+S04+S03)$ | 725.71 | 1.56 | 0.076 | 0.72 |
| $P(ROAD)\psi(S05+HUVe+S04)$ | 725.78 | 1.63 | 0.074 | 0.79 |
| $P(ROAD)\psi(S05+HUVe+SED+S04+LIV)$ | 725.80 | 1.65 | 0.073 | 0.86 |
| $P(ROAD)\psi(S05+HUVe+SED+S04+ELE)$ | 725.83 | 1.68 | 0.072 | 0.93 |
| $P(ROAD)\psi(S05+HUVe+SED+RD)$ | 725.97 | 1.81 | 0.067 | 1.00 |
| $M_m$ | | | | |
| $P(ROAD)\psi(T00+S05+SED+S04+RD+HOME)$ | 584.28 | 0.00 | 0.67 | 0.67 |
| $P(ROAD)\psi(T00+S05+SED+S04+RD+HOME+ELE)$ | 585.68 | 1.40 | 0.33 | 1.00 |

Note: $\Delta AIC$ is the difference in the Akaike information criterion (AIC); AIC $W_t$ is the model weight; and Cum. $W_t$ is the cumulative weight. $P$ : probability of detection; $\psi$ : probability of occupancy. A detailed description of the variables is provided in Table 1.

### Detection probability and occupancy probability estimation

The tiger detection probabilities (*P*) across the study area were 0.18 (SE = 0.01) and 0.16 (SE = 0.01) for M_0_ and M_m_, respectively, and there was no significant difference between M_0_ and M_m_ (Table 3). No significant differences were observed between the models when the detection probabilities of camera trapping sites inside and outside of the home range area were considered (Table 3). We found that tiger detection probabilities were higher inside compared with outside their home ranges (Table 3).

**Table 3.**
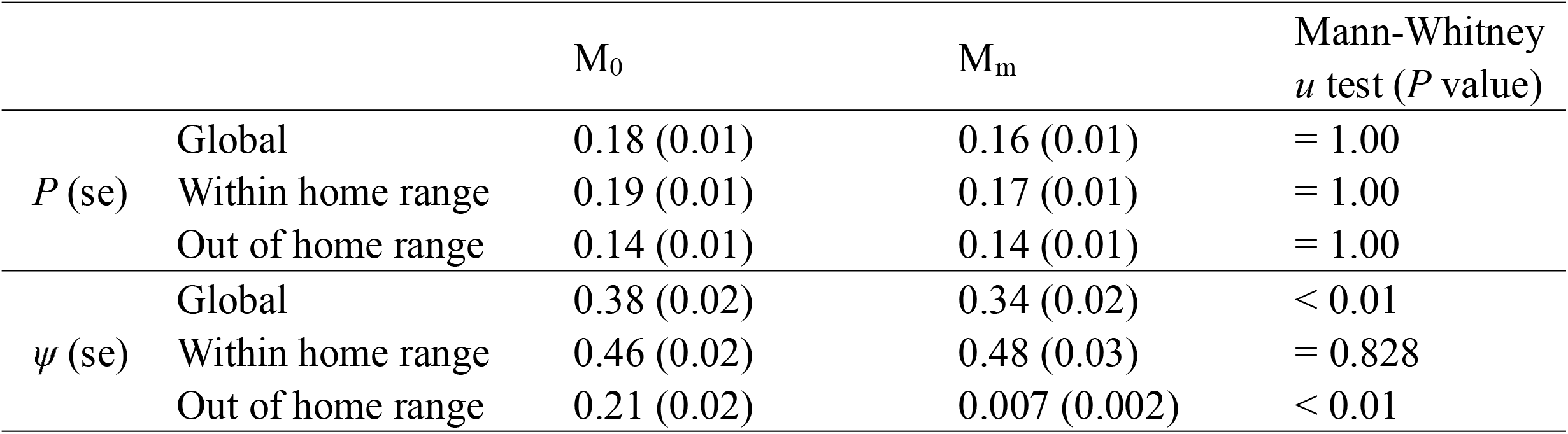
Probability of detection (*P*) and occupancy (ψ) from M_0_ and M_m_. The Mann–Whitney *u* test was used to test the differences between the two processing methods.

Occupancy probabilities (ψ) differed significantly between M_m_ and M_0_ (Table 3, Fig. 4). The global occupancy probabilities for M_m_ (0.34 ± 0.02) were slightly but significantly lower than those for M_0_ (0.38 ± 0.02) (Table 3). Although there were no significant differences in the occupancy probability in home ranges (0.48 ± 0.03 for M_m_ and 0.46 ± 0.02 for M_0_), they were 30 times lower for M_m_ (0.007 ± 0.002) than for M_0_ (0.21 ± 0.02) outside home ranges (Table 3), which was supported by the occupancy probability map (Fig. 4).

**Figure 4.**
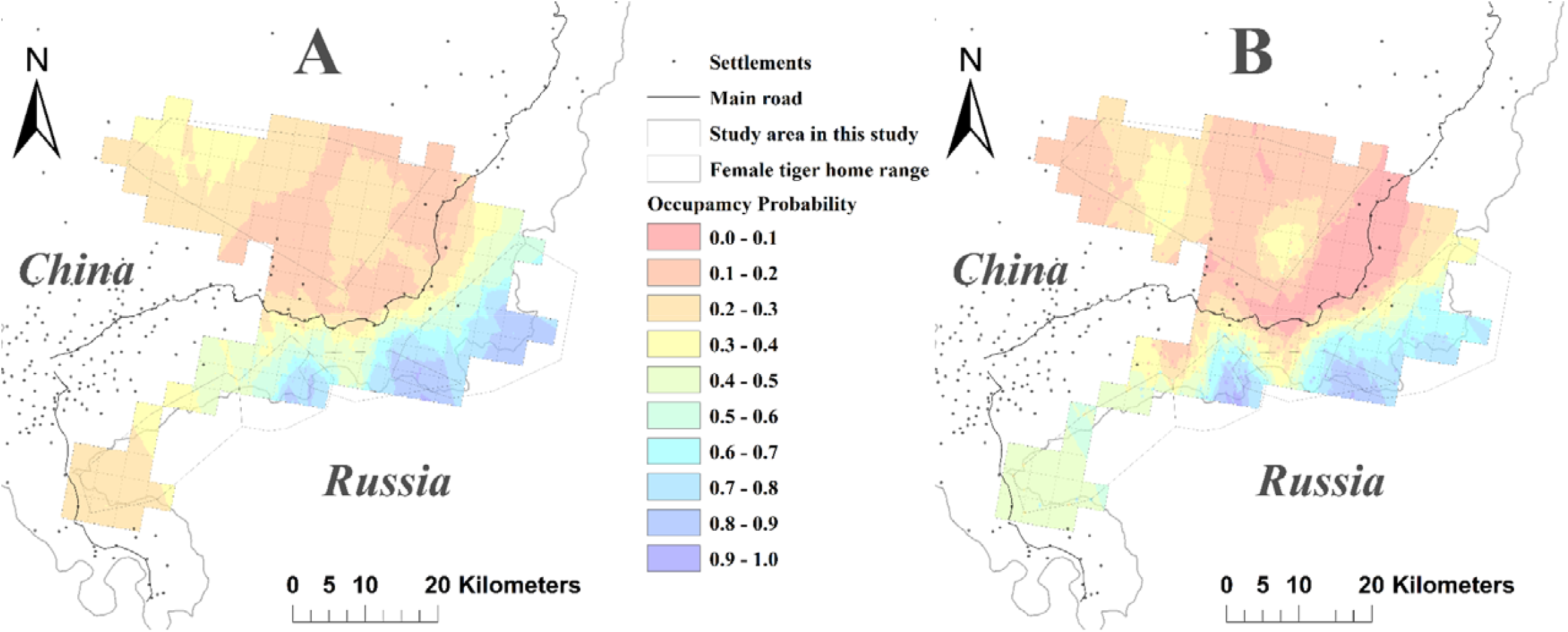
Map of the probability of occupancy from the occupancy model in the study area. A is the result from M_0_; B is the result from M_m_.

### Effects of variables on detection and occupancy estimation

The Z scores of all variables ranged from -1.96 to 1.96 and showed good convergence (Table 4). The detection probability of tigers was only significantly affected by dirt roads in the conventional and modified occupancy models (Tables 2, 4).

**Table 4.** Variable estimates and 95% credible intervals (CIs) from M_0_ and M_m_. Estimates of coefficients are reported for standardized variables, scaled to the mean and standard error (SE). Bold face indicates that variables had a significant association with Amur tiger habitat use and detection because their 95% CIs did not overlap with zero. |Z| < 1.96 indicates model convergence.

|  | Variables | Mean | SE | 95% CI | Z score |
| --- | --- | --- | --- | --- | --- |
| $M_0$ | | | | | |
| Detection<br>($P$) | Intercept | **-1.46** | **0.31** | **(-1.96, -0.95)** | **1.17** |
|  | Road type (Dirt road) | **1.29** | **0.32** | **(0.77, 1.82)** | **-0.72** |
|  | Road type (Forest trail) | 0.46 | 0.34 | (-0.11, 1.02) | -0.11 |
|  | Road type (Logging road) | 0.33 | 0.35 | (-0.25, 0.90) | -0.83 |
|  | Road type (Ridge) | 0.16 | 0.35 | (-0.40, 0.76) | -1.82 |
| Habitat use<br>($\psi$) | Intercept | -0.12 | 0.25 | (-0.51, 0.28) | -0.24 |
|  | **S05** | **1.53** | **0.52** | **(0.70, 2.31)** | **-0.67** |
|  | **HUVE** | **0.29** | **0.14** | **(0.06, 0.50)** | **-0.51** |
|  | **SED** | **0.32** | **0.17** | **(0.04, 0.58)** | **1.33** |
|  | S04 | -0.35 | 0.22 | (-0.71, 0.01) | -0.38 |
| $M_m$ | | | | | |
| Detection<br>($P$) | (Intercept) | **-1.56** | **0.36** | **(-2.15, -0.97)** | **-0.40** |
|  | Road type (Dirt road) | **1.09** | **0.37** | **(0.49, 1.71)** | **0.40** |
|  | Road type (Forest trail) | 0.30 | 0.40 | (-0.36, 0.94) | -0.18 |
|  | Road type (Logging road) | 0.51 | 0.40 | (-0.15, 1.17) | -0.07 |
|  | Road type (Ridge) | 0.33 | 0.40 | (-0.33, 0.98) | 0.11 |
| Habitat use<br>($\psi$) | **(Intercept)** | **-2.90** | **1.17** | **(-4.75, -1.09)** | **0.30** |
|  | **T00** | **2.01** | **0.93** | **(0.70, 3.29)** | **0.31** |
|  | **HOME** | **3.24** | **1.20** | **(1.35, 5.09)** | **-0.15** |
|  | **S05** | **1.00** | **0.61** | **(0.05, 1.85)** | **0.23** |
|  | **RD** | **0.40** | **0.22** | **(0.03, 0.75)** | **1.67** |
|  | SED | 0.01 | 0.20 | (-0.32, 0.34) | 1.25 |
|  | S04 | -0.37 | 0.27 | (-0.80, 0.04) | 0.19 |

Note: A detailed description of the variables is provided in Table 1.

In M_0_, S05, HUVE, and SED had significant positive effects on the occupancy models (*P* < 0.05), and S04 had a negative effect (*P* > 0.05) (Table 4). The effects of SED, S05, and S04 in M_m_ were similar to those in M_0_; however, the effect of SED was not significant, and HUVE was not an important variable in M_m_ (Tables 2, 4, and S3). The M_m_ results indicated that T00, HOME, and RD had significant positive effects on the occupancy probability of tigers (*P* < 0.05) (Tables 2, 4, and S3). Maps based on occupancy models showed that areas with higher occupancy probability were concentrated in the home ranges of tigers, especially home ranges in the boundary area (Fig. 4). We also found that lower occupancy probability was concentrated near the main road and its surroundings in M_m_, which was in contrast to the M_0_ map (Fig. 4).

## Discussion

Accurate characterization of the relationship between the distribution of wildlife species and environmental factors at the landscape scale is critically important for the conservation of large solitary carnivores (Karanth *et al*. 2011). We examined the effect of ‘True-Presence’ and ‘Pseudo-Presence’ associated with the territorial behaviour of Amur tigers on the performance of occupancy models. Our results indicated that M_m_, which considered the ‘True-Presence’ and ‘Pseudo-Presence’ in the models, was more accurate, and the results of this model could more effectively explain the distribution and habitat use of the Amur tigers in our study area. By contrast, M_0_ significantly overestimated the probability of occupancy outside of the home range area, as detection probabilities did not significantly differ between M_0_ and M_m_. We also acquired more reasonable variables on the habitat selection of the Amur tiger from M_m_, which could facilitate the development of conservation policies for the tigers in our study area.

### Performance assessment of M_0_ and M_m_

Although all indicators of the performance of both M_0_ and M_m_ indicated no lack of fit of our models, all goodness-of-fit values for M_m_ were superior to those for M_0_ (Tables 2 and S3–S4). The number of competing models of M_0_ was greater than that of M_m_ (11 vs. 2), and the AIC of the top model from M_0_ was much higher than the AIC of the top model from M_m_ (724.15 vs. 584.28); the cumulative weight of the top model from M_0_ was much lower than that of the top model from M_m_ (0.17 vs. 0.67). All these results indicate that M_m_, which was based on fewer variables, was more powerful for explaining the effects of environmental factors on the habitat use of Amur tigers (9 and 8 variables for M_0_ and M_m_, respectively) (Tables 2, 3). Therefore, M_m_ permits more reasonable characterizations of the habitat use of the Amur tiger (from a statistical perspective).

### Occupancy probability

Local tiger occupancy is strongly determined by prey density and human disturbance (Smith 1987; Goodrich *et al*. 2010; Karanth *et al*. 2011; Yang 2018). Sika deer significantly affected the occurrence of the Amur tiger in both M_0_ and M_m_ (Table 4). Our results indicate that the densities of medium to large ungulate prey species are determinants of tiger presence at a fine scale (Karanth *et al*. 2010; Yang *et al*. 2018). Thus, the higher occupancy probabilities of tigers concentrated along Sino-Russia could be partially explained by the distribution of sika deer along the border (Wang *et al*. 2016) (Fig. 4). Roe deer was relatively important in both M_0_ and M_m_ but had no significant effect on the occupancy probability of Amur tigers (Tables 4 and S3). Tigers tend to avoid preying on roe deer because of their small body size (Andheria, Karanth & Kumar 2007; Yang *et al*. 2018).

It is not surprising that prey species have the same impacts on tiger occupancy. However, it is indisputable that human disturbances mediate tiger distribution, which determining the ‘True-Presence’ and ‘Pseudo-Presence’ of tigers. In M_0_, the HUVE and SEM have significant opposite impacts on occupancy probability, which are difficult and illogical for explaining the influences of HUVE and SEM on tiger occupancy from an ecological aspect (Table 4). Human activity and settlements have been confirmed to have negative effects on big cats, through prey depletion, direct poaching and habitat encroachment (Carroll & Miquelle 2006; Barber-Meyer *et al*. 2013; Wang *et al*. 2016), which are detrimental to tiger habitat use. In M_m_, RD instead of human activity, showed a significant impact on tiger occupancy, indicating that tiger tends to avoid the main road in our study area (Table 4, Fig. 4). Roads have already been reported to reduce Amur tiger survival rates, due to the poaching of tigers and their prey as well as collisions with vehicles (Chapron *et al*. 2008; Goodrich *et al*. 2008). Wang *et al*. (2016) indicated that a road-cultivation zone consisting of the main road (running parallel to the border), farmland, and settlements cuts across the region and hinders the dispersal of tigers and their prey into China. Smith (1984) suggested that tigers avoided farmland during dispersal. We found that the home ranges of tigers tended to be located far away from the main roads (Fig. 3). Maps of land cover and the probabilities of occupancy from the occupancy model in the study area also indicated that the area around the main road is not suitable for tigers (Figs. 1, 4). The probability of occupancy outside the home range area from M_0_ is approximately 30 times higher than that from M_m_, which indicates that consideration of ‘True-Presence’ and ‘Pseudo-Presence’ provides more accurate estimates of the habitat use of tigers compared with M_0_ (Table 4). The sporadic presence in an unsuitable area for tigers is denoted as ‘Pseudo-Presence’ in M_0_, which does not reflect actual occupancy or habitat use. The more ecologically meaningful results from M_m_ fully prove that the territorial behaviour should be fully considered when studying habitat selection or distribution estimation of tigers through occupancy models (Tables 3-4, Figs. 3-4).

### Conclusion

As a method to study the influence of territorial behaviour on occupancy model performance, this study is very meaningful for accurate habitat selection and distribution estimation, which is more conducive to assessing the distribution status of wildlife. Our method could be applied to other species with territorial behaviour, which could enhance the accuracy of occupancy models in habitat selection and distribution estimation for wildlife.

## Authors’ contributions

Haitao Yang and Limin Feng conceived the ideas and designed methodology; Haitao Yang, Limin Feng collected the data; Haitao Yang, Bing Xie, Yinan Gong, Yanwen Fu analysed the data; Yinan Gong printed the figures; Bing Xie and Haitao Yang led the writing of the manuscript. All authors contributed critically to the drafts and gave final approval for publication.

## Acknowledgements

We gratefully acknowledge the Chinese National Forestry and Grassland Administration, Northeast Tiger and Leopard National Park, and all rangers with their fieldwork for their assistance in field sampling, measurements, and data input. We also thank the hard and outstanding fieldwork of the Land of the Leopard National Park (LLNP) in Russia’s Southwest Primorye Province related to data of the tiger’s home range estimation.

## Declaration

The authors declare no conflicts of interest.

## Notes

### Competing Interest Statement

The authors have declared no competing interest.

