## Appendix 1-3 and Table S1-S4 for "Does the territorial behaviour of the Amur tiger affect the accuracy of occupancy estimation?": Appendix 2.docx

**Materials and methods**

*Study area*

The occupancy models were only used the data from the Chinese side, the area is located in the northern portion of Changbai Mountain in the Northeast Tiger and Leopard National Park (NTLNP), Jilin province, Northeast China, bordering Southwest Primorye, Russia to East and North Korea to the southwest (Fig. 1). This region’s Amur tiger population is thought to be the source population for the Amur tiger recovery in China. As a result, this area is also considered the highest priority Tiger Conservation Area in China (Hebblewhite *et al.* 2012; Wang *et al.* 2016). The altitude ranges from 5 to 1,477 m. The climate is characterized as a temperate continental monsoon with a frost-free period of 110 - 160 days/year. The average temperatures range from 3.90 - 5.65 ℃, and the average precipitation is 580 - 618 mm with the most precipitation occurring from June to August. With the implementation of the Natural Forest Conservation Program (NFCP) and Grain-to-Green Program (GTGP), the commercial logging of natural forests has been halted. The main commercial activities in the rural areas are free-range cattle grazing, edible ferns, ginseng farms, and frog farming.

*Modified occupancy model*

We ran models in two phases: 1) models without spatial autocorrelation to select covariates and 2) models with spatial autocorrelation. The occupancy models for the conventional and modified occupancy models were modeled separately.

The first phase was performed under a maximum likelihood framework and conducted with the R package unmarked version 0.12-2 (Fiske *et al.* 2015). First, we modeled the detection probability (*p*) by using all combinations of covariates (Table 1) while holding *ψ* constant, and the top models with ∆AIC < 2 were considered to contribute equally and used to model habitat use probability in relation to the site covariates (Tan *et al.* 2017). Then, we ran each combination of site-specific covariates while the detection probabilities were modeled following models selected previously. The top models with ∆AIC < 2 were considered to contribute equally. We selected only covariates with summed model weights > 0.5 (Kalies *et al.* 2012) to model spatial autocorrelation in the next phase. We assessed the goodness-of-fit of the global model to evaluate the probability that the model would be correct (*P* > 0.5) and the accuracy of estimation determined by c-hat (0.5 < c-hat < 1.5) (MacKenzie & Bailey 2004; MacKenzie *et al.* 2017).

In the next phase, the R package *stocc* version 1.30 (Johnson *et al.* 2013) was used to model spatial autocorrelation with restricted spatial regression (RSR). The posterior predictive loss criterion (PPLC) was used to compare models without spatial autocorrelation parameters (top-ranked models from the first phase) to models with spatial autocorrelation (Bayesian RSR) (Gelfand & Ghosh 1998). The moran.cut cut-off parameter was set to 20 (Goodrich *et al.* 2010; Hughes & Haran 2013). We set flat prior distributions for *p* and *ψ* and a gamma (0.5, 0.00005) distribution for the spatial component (Johnson *et al.* 2013). We ran the Gibbs sampler for 10,000 iterations, with a burn-in of 1,000 iterations, to estimate the parameter mean, standard deviation (SD), and 95% Bayesian credible interval (CI). Covariates with a 95% CI that did not overlap with 0 were considered to have a significant association with tiger detection and habitat use. We used Geweke diagnostic statistics (Geweke 1991) to assess model convergence (|Z| < 1.96).
