## Appendix 1-3 and Table S1-S4 for "Does the territorial behaviour of the Amur tiger affect the accuracy of occupancy estimation?": Appendix Table S1.docx

Table S1 Estimation of the female tigers’ home range size based on the 100% minimum convex polygons (MCP)

| Tiger ID | Sample period | Home range size km^2^ (100%MCP) |
| --- | --- | --- |
| TIG-03 | 2011.06 - 2015.05 | 116.52 |
| TIG-04 | 2012.07 - 2015.04 | 329.99 |
| TIG-05 | 2014.04 - 2015.10 | 528.83 |
| TIG-09 | 2012.11 - 2016.04 | 211.84 |
| TIG-22 | 2012.10 - 2016.04 | 102.59 |
