## Appendix 1-3 and Table S1-S4 for "Does the territorial behaviour of the Amur tiger affect the accuracy of occupancy estimation?": Appendix Table S2.docx

Table S2 Correlated variables among all covariates

|  | T00 | ELE | RD | SED | LIV | HUVE | S03 | S04 | S05 |
| --- | --- | --- | --- | --- | --- | --- | --- | --- | --- |
| T00 | 1.00 | -0.17 | -0.04 | 0.10 | -0.10 | 0.26 | -0.08 | -0.16 | 0.30 |
| ELE |  | 1.00 | 0.17 | 0.53 | -0.15 | -0.25 | 0.26 | 0.44 | -0.09 |
| RD |  |  | 1.00 | 0.18 | 0.01 | -0.17 | 0.02 | 0.09 | -0.11 |
| SED |  |  |  | 1.00 | -0.17 | -0.05 | 0.13 | 0.17 | 0.16 |
| LIV |  |  |  |  | 1.00 | 0.24 | -0.09 | -0.18 | -0.16 |
| HUVE |  |  |  |  |  | 1.00 | -0.13 | -0.23 | 0.04 |
| S03 |  |  |  |  |  |  | 1.00 | 0.30 | 0.21 |
| S04 |  |  |  |  |  |  |  | 1.00 | -0.14 |
| S05 |  |  |  |  |  |  |  |  | 1.00 |

Note: Detailed description of variables is in Table 1.
