## Appendix 1-3 and Table S1-S4 for "Does the territorial behaviour of the Amur tiger affect the accuracy of occupancy estimation?": Appendix Table S3.docx

Table S3 Summary of summed model weights for parameters derived from the Top-ranked occupancy models, see Table 2

|  | Variables | Summed model weights |
| --- | --- | --- |
| M_0_ | | |
| Detection (*P*) | Road type | 1.00 |
| Habitat use (*ψ*) | Sika deer | 1.00 |
|  | Human activity | 1.00 |
|  | Settlement distance | 0.85 |
|  | Roe deer | 0.54 |
|  | Wild boar | 0.17 |
|  | Elevation | 0.16 |
|  | Road distance | 0.15 |
|  | Livestock | 0.07 |
| M_m_ | | |
| Detection (*P*) | Road type | 1.00 |
| Habitat use (*ψ*) | Non-settled tigers and males | 1.00 |
|  | Sika deer | 1.00 |
|  | Settlement distance | 1.00 |
|  | Roe deer | 1.00 |
|  | Road distance | 1.00 |
|  | Home range | 1.00 |
|  | Elevation | 0.33 |

Note: *P*: the probability of detection; *ψ*: the probability of occupancy. A detailed description of variables is in Table 1.
