## Appendix 1-3 and Table S1-S4 for "Does the territorial behaviour of the Amur tiger affect the accuracy of occupancy estimation?": Appendix Table S4.docx

Table S4 Spatial autocorrelation analysis of occupancy model without considering individual tiger home range and with considering individual tiger home range.

|  | Model | M_0_ | M_m_ |
| --- | --- | --- | --- |
| D.m | None | 215.42 | 166.56 |
|  | RSR | 215.50 | 166.26 |
| G.m | None | 106.30 | 81.35 |
|  | RSR | 106.28 | 81.08 |
| P.m | None | 109.13 | 85.21 |
|  | RSR | 109.21 | 85.18 |

Note: D.m: Goodness of fit + Complexity penalty; G.m: Goodness of fit; P.m: Complexity penalty; None: without considering the spatial autocorrelation; RSR: Restricted Spatial Regression (with considering the spatial autocorrelation).
